# The evolution of fitness during range expansions in different dimensions

**DOI:** 10.1101/2023.12.29.573608

**Authors:** Kotsar Yurii, Hikaru Matsuoka, Gen Tamiya

## Abstract

We develop a set of programs – the range expansions simulation kit (RESK) – to efficiently simulate range expansions of populations on a square lattice in 1D, 2D (on a strip, and on a disk) and 3D (in a cylinder, and in a sphere). In this study, we present the simulation kit to the public and present some results of using it to simulate a population of diploid individuals with finite genome regions, each containing infinite sites. Specifically, using the programs, we calculate and analyse the temporal evolution of population fitness, as well as fitness on the expansion front, in 3D for the first time. We explore the model over different conditions, compare normalisation methods for fitness, and explore the case of radial (sphere) and axial (cylinder) expansions in 3D, which might apply in the analysis of the different real-life populations, such as viruses/bacteria inside a host, marine species in ocean environments, and potentially in future space colonization planning. In 3D expansions, we find complex spatial fluctuations in deme-average fitness values, different from those in radial 2D expansions. In axial 3D (cylinder) expansions, we determine that the highest-valued deme-average fitness lies along the axis of the expansion. We also find the fluctuation patterns of fitness in 3D cylinder expansions, similar to those previously seen in radial expansions in 2D. In radial 2D (disk) expansions, we find that the fitness of a population undergoing multiple mutations shows a smooth combination of binary segregation pictures against each of those mutations. We confirm the accumulation of deleterious mutations -- a phenomenon known as expansion load -- in all scenarios above. We present the software used in the above to the public as a Julia repository, ready to use with different functions for simulating range expansions with minimal syntax.

## 1. INTRODUCTION

Most species on Earth exhibit fluctuating habitats, a significant dynamic mode of which is range expansions. Understanding the consequences and dynamics of such changes of habitats is of great importance in evolutionary biology. Their distribution, however, is nontrivial even in a spatially deterministic model of migration, since almost all organisms have multiple segregating genetic loci which accept random mutations (which, in turn, affect the fitness, and thus, the number of individuals). Due to demes on the expansion front having less genetic variation in the early generations after colonisation, while the front is moving, a mutation (neutral, deleterious or beneficial) can fix itself on the front for many generations thereafter, but also spread back into the bulk of the population. This is the principle of *gene surfing* (Klopfstein *et al*., 2005). Through many generations, the effect of those mutations will accumulate, if the mutations are prevalently either beneficial or deleterious (Peischl *et al*., 2013). There is evidence to show that 70% to 90% of all mutations are deleterious (Eyre-Walker and Keightley, 2007).

This lowers the value of fitness averaged over the individuals in every deme on the expansion front – simply “*mean front fitness*” hereafter – and exacerbates the difference in fitness between the core population and the expansion front. Nevertheless, if the selection is soft, mean front fitness can never fall to zero, and a population can avoid extinction (Peischl *et al*., 2015a). In this regime, establishing the means to quantify the change in mean front fitness over the expansion front is an important goal, the results of which have various applications in the field of population ecology, especially in the wake of some range expansions caused by climate change (Bokma, 2010; Parmesan, 2006; Pateman *et al*., 2012; Yamano *et al*., 2011). These cases assume a one-dimensional (1D) or two-dimensional (2D) habitat, but this change in fitness could also be modelled for three dimensions (3D), which is the focus of this study. In the 3D case, quantifying the change in mean front fitness is important in elucidating the capabilities of viruses (Bocharov *et al*., 2016), as well as marine species to spread (Kitchel *et al*., 2022; Assis *et al*., 2016), or could be vital information in the planning stages of human space expansion. A particularly important case is the gradual migration of marine species in depth, in addition to lateral migration – primarily to deeper latitudes, as a result of climate change (Poloczanska *et al*., 2016). In these scenarios, a reliable way to simulate and understand range expansions in three dimensions is required, especially for the case of an expansion along an axis and a certain radius around it to allow for slight deviations -- in our terminology, the *3D cylinder* scenario.

So far, an analytical approximation to the change in mean front fitness has been constructed only for 1D (Peischl *et al*., 2013). This prior research shows that it’s difficult to construct a theoretical formula for mean front fitness for the case of range expansions with logistic growth, mutation, selection and genetic drift, and so it is valuable to create a set of simulation programs with the purpose of generating the dynamics of a range expansion, generation by generation, and to use the simulated data to quantify the change in mean front fitness. We thus set the goal of developing a time-effective and robust in terms of starting conditions simulation kit, that can also work with 2D and 3D expansions. Furthermore, with this study, we set out to provide an example of our simulation kit with some realistic parameters for human range expansions.

We base this study in part on the research in Peischl *et al*. (2013). Their study includes a C++ program, which is able to generate fitness values for each deme in a one-dimensional expansion. In their publication, some simulation results for 1D and 2D have been given. In our study, based on the aforementioned C++ program, as well as Gilbert *et al*. (2018), we recreate the functionality for 1D expansions, and independently develop 2D and 3D functionality, in the Julia programming language. In this study, we consolidate the above into a ready-to-use simulation kit and present it as well as examples of simulations in different dimensions, while exploring different steps throughout the algorithm (such as normalisation methods). It is available in the “Data availability statement”.

## 2. METHODS

### 2.1.Model: genome regions with infinite sites

Our model is based largely on the model in Peischl *et al*. (2013). For all dimensions, we start with a habitat of discrete laterally connected lattice of demes, and employ a model which accepts a population of monoecious diploid individuals of size *N*, employing the stepping-stone model (Kimura, 1953), where migration is only present between adjacent demes.

In this model, each diploid individual has a genome of 2*L* independently segregating regions with no epistasis and no dominance. We set *L* = 20 in this study. These regions are the basic units during genetic recombination, which occurs once per generation. During genetic recombination, regions 1 ≤ *k* ≤ *L* have a 1/2 chance toswap with their pair in *L+*1 ≤ *k* ≤ *2L*. Inside these genome regions, we assume the infinite-site model, so the same segregating region could have multiple mutations. A mutationhas a fixed chance *μ* of occurring in a gamete, and upon that, a fixed chance ϕ of being deleterious. We set this value as ϕ = 0.9, unless stated otherwise. Deleterious; beneficial mutations change the fitness of a recombining region by 1 - |*s*| ; 1 + |*s*| respectively. In this study, |*s*| is a fixed value in each simulation trial, and is called the *selection coefficient*. Multiplicative fitness is assumed. The fitness values are the main variables in our model; they are stored for every segregating region for every individual and begin at 1. In order to model mutations, we sample the number of mutations from a Poisson distribution with mean *μ*, determine whether the mutation is beneficial or deleterious, then act on the fitness of the recombining region picked randomlyfrom a uniform distribution. Since an infinite-site model is assumed inside the genome regions, the chance of further changing the allele of the samelocus is infinitesimal, thus once the mutation has affected the fitness, the multiplicative factor of its selective effect always persists in the individual fitness. The evolution of each deme’s population is based on the Beverton-Holt formula for discrete logistic growth:

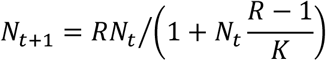

where *N*_*t*_ is the number of individuals in a deme in generation *t, R* = 2 is the proliferation rate, and *K* is the constant carrying capacity, set equal to 100, unless specified otherwise.

Furthermore, we sample the actual number of offspring individuals from a Poisson distribution with mean *N*_*t* +1_. The deme itself does not have a hard limit for the number of individuals, hence due to Poisson sampling, the number of individuals in a deme can sometimes exceed *K*. The fitness of all the segregating regions is multiplied for all individuals within a deme, and pairs of individuals are randomly selected from the deme. The maximum fitness within a deme is found, and the individuals from pairs have their fitness divided by the maximum fitness and compared with a uniform random variable. If both individuals pass, their zygote enters into the new generation. This is done until the number of offspring sampled above is satisfied. Since we assume monoecious individuals, selfing is allowed.

Finally, each generation, migration is allowed and equally occurs between all adjacent demes, with a total migration rate *m*. In one dimension, migration rate is simply *m*/2 to move to either of two adjacent demes. In higher dimensions, migration rate is *m*/*l*, where *l* is the number of laterally adjacent demes. In particular, l is 2 in 1D, 4 in 2D, and 6 in 3D. We then employ an additional check for the demes at the boundaries of the habitat: we use reflecting borders, in the sense that if migration has been initiated, the individual shall always move somewhere -- in this case, away from the boundary of the habitat.

It is difficult, in principle, to calculate a clear proportion of deleterious and beneficial mutations, since accurate measurement of the effects of single mutations is possible only when they have fairly large effects on fitness (> 1%) (Eyre-Walker and Keightley, 2007). Nevertheless, different studies are in agreement that the proportion of deleterious mutations is prevalent and is between 70% and 90% (Kim *et al*., 2017). In our model, single mutations can only have constant deleterious or beneficial effects, which we set for this study as *s* =−0.005 and *s* =+0.005 respectively, and which can be viewed as average values of all possible mutations for −1 < *s* < 0 and 0 ≦*s* < ∞ respectively, where the weights for *s* =−0.005 and *s* =+0.005 (multiplicative changes in fitness of 1 - 0.005, 1 + 0.005 respectively) need to be decided. We base the weights on the distribution values in humans (Eyre-Walker and Keightley, 2007): ∼ 90% for deleterious and ∼10% for beneficial mutations, when taken without neutral mutations. The proportion of deleterious to all mutations can be changed in the program, and our model can be used for viruses and other populations as well: Sanjuan *et al*. (2010) give ∼ 70% for deleterious mutations in viruses. There are also lethal mutations, which have to be considered due to their special logic of potentially erasing an individual from the model. As in the current study we are focusing on human populations, for which lethal mutations make up less than 0.5% of deleterious mutations (Gao *et al*., 2015), we can reasonably ignore them. For viruses, however, more than 25% of mutations could be lethal (Sanjuan *et al*., 2010). A more accurate approach would be to consider variable *s* (e.g. multiple *s* bands), which we aim to integrate in our simulations in the future.

### 2.2. Simulations

We developed multiple methods for our simulations kit to recreate different scenarios of range expansions, based on the model described above. These methods are named *rangeexp_#d_inf*, where # is the number of spatial dimensions.

We divide the total number of generations into a burn-in phase and an expansion phase. During burn-in, the habitat is restricted to an area much smaller than the full space. This is done to allow the population to reach mutation-selection-drift equilibrium (Peischl *et al*., 2013). During the expansion phase, the population is allowed to migrate freely, up until the boundaries of the habitat. In all simulation trials (1D, 2D and 3D), we picked a size of the habitat that allow the population to fill the habitat up in several hundreds to thousands of generations. Next, we describe the general algorithm of the programs (Figure 1).

**FIGURE 1.**
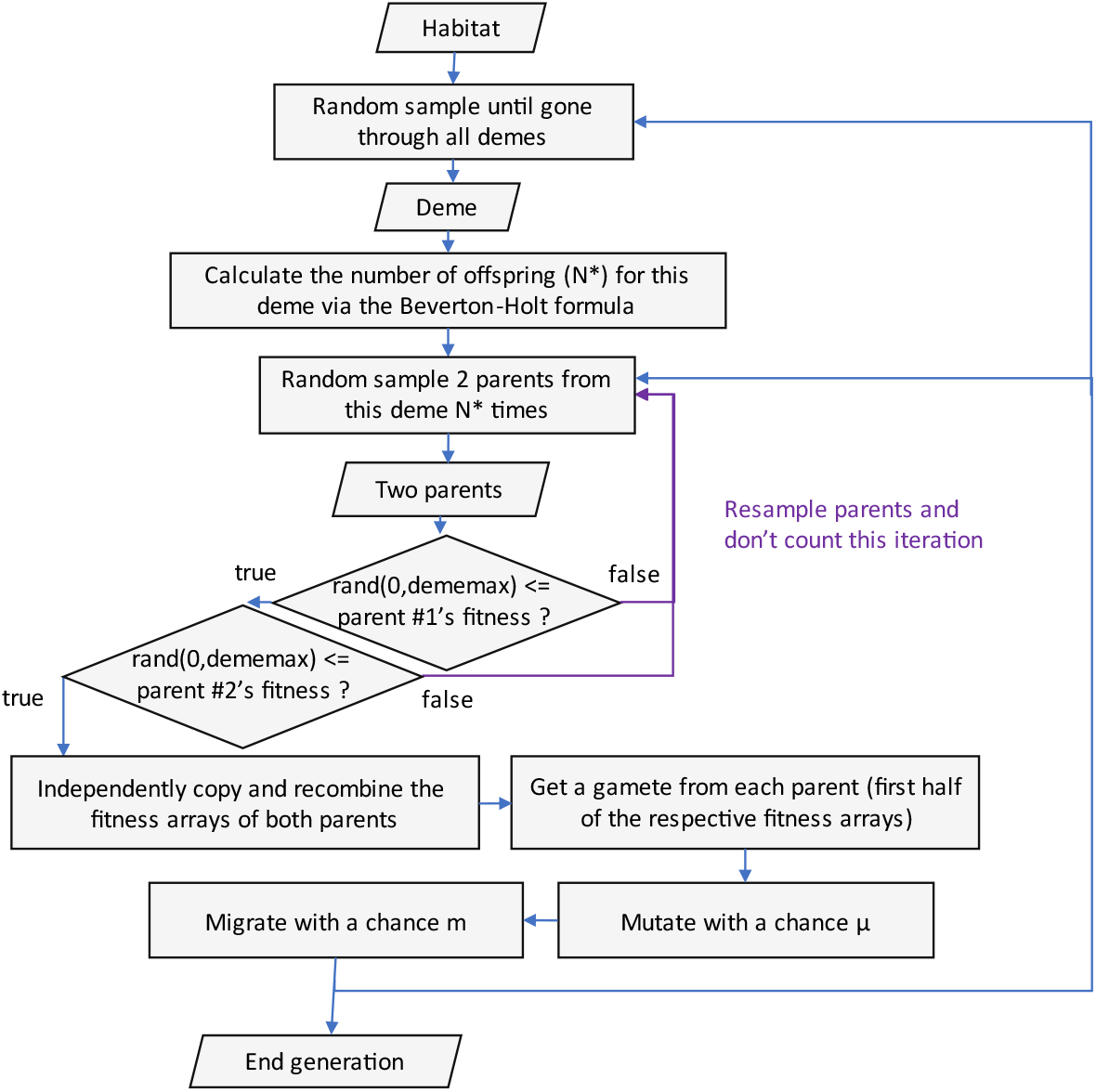
The algorithm of the cycle in each generation of expansion in our program. “dememax” is the maximum fitness in the deme.

We start with an array of empty demes, which are themselves arrays of individuals (Figure S1). We then pick *b* random demes and fill them to their carrying capacity (given in Table 1) with individuals. Every individual starts out with an array of fitness values of 1 for every genome region. For all simulations in the current study, we set this parameter *b* to 5. Every generation, we run a cycle over all demes: we first determine the number of offspring logistically (as described above), then, for every individual in the offspring, we iterate over each individual in the current generation, in a random order, until we find suitable parents for the offspring individual (we call this *draw until pass* below). Specifically, the two candidates for parents must draw samples from a uniform distribution from 0 to the current deme’s maximum fitness, that pass the fitness values of their gametes. In one case below (referenced as *first pair*), we also employ a variation wherein, instead of a running a *while* cycle, we run a *for* cycle on the parents: in other words, the fate of offsprings individuals depends on the first random pair of parents to either pass or fail the check; if this check fails, the offspring is automatically less by one individual.

**TABLE 1.** List of simulation parameters and their values used in this paper.

| Parameter | 1D (1) | 1D (2) | 1D (3) | 2D strip | 2D disk | 3D cylinder | 3D sphere |
| --- | --- | --- | --- | --- | --- | --- | --- |
| Number of replicates | 70 | 10 |  | 20 |  | 10 |  |
| Burn-in generations | 10000 | 5000 |  | 1000 | 1000 | 1000 | 1000 |
| Expansion generations | 2500 | 1250 |  | 1000 | 500 | 1000 | 150 |
| Space size (burn-in) | $1 \leq X \leq 5$ | | | $1 \leq X \leq 5,$<br>$1 \leq Y \leq 10$ | $0 \leq R \leq 3$ | $1 \leq Z \leq 5,$<br>$0 \leq R \leq 10$ | $0 \leq R \leq 3$ |
| Space size | $1 \leq X \leq 500$ | | | $1 \leq X \leq 250,$<br>$1 \leq Y \leq 10$ | $0 \leq R \leq 100$ | $1 \leq Z \leq 300,$<br>$0 \leq R \leq 10$ | $0 \leq R \leq 25$ |
| Selection coefficient | 0.005 |  | 0.03 | 0.005 |  |  |  |
| Mutation rate | 0.05 |  | 0.1 | 0.05 |  |  |  |
| Prop. of del. mutations | 0.90 | 0.45 | 0.90 |  |  |  |  |
| Migration rate | 0.05 |  |  |  |  |  |  |
| Carrying capacity | 100 |  |  |  |  |  |  |
| Filled demes at start | 5 |  |  |  |  |  |  |
| Proliferation rate | 2 |  |  |  |  |  |  |
| Number of gene regions | 20 |  |  |  |  |  |  |

After this step, the two parents’ gametes are recombined, and mutations are introduced. The offspring individual is formed, and then allowed to migrate.

The are several parameters that the simulations take as input. We summarise them in Table 1. The simulations output:

- the **deme-average fitness matrix** per every generation, which stores the deme-average individual fitness for every deme in the habitat,
- the **population matrix** per every generation, which stores deme populations,
- the **deme-average mutations matrices** per every generation, storing the deme-average number of deleterious as well as beneficial mutations.

The focus for this study is to use the output for the deme-average fitness to further calculate its average over the expansion front, and then graph the temporal evolution of the resulting mean front fitness. We do this for different dimensions and configurations, and compare it to the analytical approximation found below in 2.3. More specifically, we find the expansion front as the collection of outermost demes, counting from the borders of the habitat to one side (*x* > 0 half-plane in 2D, and *z* > 0 half-space in 3D) in axial expansions, and all sides in radial expansions. We then find the mean of deme-average fitness values over this expansion front. The population matrix was used to confirm the logistic growth of the population, and to determine the generation at which the whole habitat was filled, i.e. the end of the expansion. The population matrix and mutations matrices are available as output options in our simulation kit.

#### 2.2.1 Normalisation of fitness

It is also common to normalise the mean front fitness. The consideration of a normalisation in time appears when we include a burn-in phase. During this phase, individuals are locked in a tight environment, and in the absence of expansion mean front fitness grows at a fast pace. It is then natural to have mean front fitness start at the same value at the onset of the expansion as the value it would have had if there was no burn-in phase. Since we set our initial value of fitness in the genome to be 1, we could normalise our mean front fitness such that the value at the onset generation is 1. To this end, we divide mean front fitness in each generation after the onset by its value at the onset of the expansion. This approach seems to yield good correspondence with data (Peischl *et al*., 2013, 2015b). We call this method “*onset mean normalisation”* in the results below.

However, this is not the only kind of normalisation applicable. A more literal definition of normalisation could also be applied, which maps the average values of fitness of each inhabited deme to the interval [0, 1] at every generation by dividing each deme-average fitness by the maximum overall inhabited demes. A similar approach is to map the values of fitnesses of each individual by dividing individual fitnesses by the maximum fitness of all surviving individuals. We have found that these approaches result in far more rapid decline in normalised mean front fitness. We included the results of dividing mean front fitness by the maximum deme-average fitness in the graphs in *Results* as “*maximum normalisation”*. Rather thanbeing a method to equate the scale of fitness values after a certain number of generations (in our case, the burn-in period) passed with the scale of fitness values obtained without that period, “maximum normalisation” is a normalisation in space, that gives a good way of quantifying the expansion load. We can also use this method to confirm that the difference in fitness between the core population and the front population grows with time in range expansions, even when the “onset mean normalisation” yields increasing fitness with time.

### 2.3 Analytical approximation

An analytical approximation to the temporal evolution of mean front fitness in this model was developed in Peischl *et al*. (2013). In this approximation, migration from a deme is only assumed possible after the deme has reached the carrying capacity *K*. This is a good approximation for low migration rates (*m*), at least compared to the logarithmic proliferation rate *r =* ln(*R*) *>> m*. Once at the capacity, *Km*/2 individuals on average migrate to further demes in one general direction (out of two). This holds especially well in axial expansions. We then only concern ourselves with those *Km*/2 individuals, since those are assumed to comprise the expansion front. We employ the Wright-fisher model of evolution of allele frequencies, but split that process into growth and sampling phases (Wahl and Gerrish, 2001). This allows us to model *T* ≈ log(2*m*^*-*1^)/*r* intermediate generations of the growth phase, during which the front population undergoes only reproduction and selection against the mutation that had just occurred. This is equivalent to a series of consequent founder effects, and is similar to a model of repeated bottlenecks.

#### 2.3.1 Probability of fixation

We derive a more general formula for the probability of fixation in this analytical model. In Peischl *et al*. (2013),the mutant frequency *x*′ in the next generation is given by

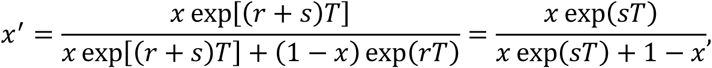

where *x* is the mutant frequency in the current generation. We would like to use the difference Δ*x* = *x*′−*x*, its mean *M[*Δ*x]* and its variance *V[*Δ*x]* , to find the probability of fixation of a mutation with effect *s* on the expansion front. We have

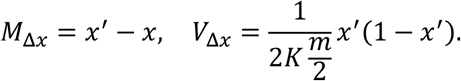

Meanwhile, the general formula for the gene frequency distribution under irreversible mutation (Kimura, 1964):

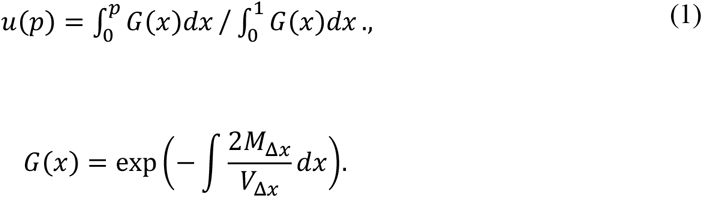

In our case:

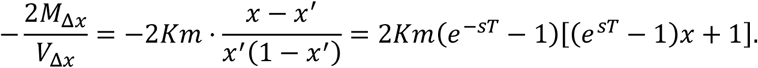

We do not assume small *s* here, and so this does not simplify further into the formula in Peischl *et al*. (2013). Next,we rearrange and define *A* and *B*:

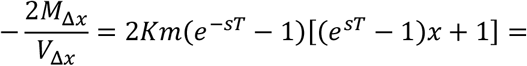

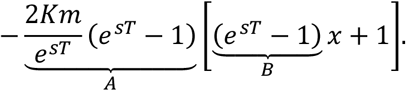

Then,

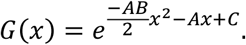

Then the integrals in Eq. 1 can be seen as Gaussian integrals. Note that *A* > 0, *B* > 0, *AB* > 0, but the coefficient of *x*^2^ in the exponent is a minus, so we are dealing with the error function erf(*x*). We have:

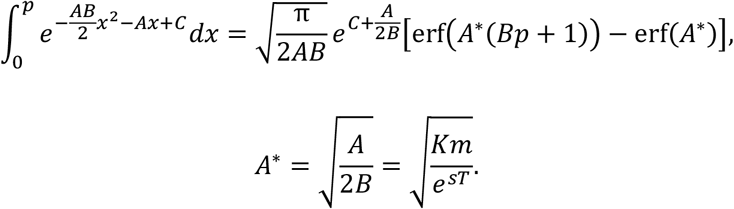

Solving the lower integral in a similar manner, Eq. 1 becomes:

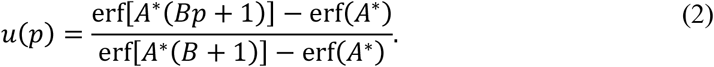

We have tested this function for the fixation probability in a flexible range of parameters *s, m, r, K*. Compared to the formula in Peischl *et al*. (2013), it gives slightly lower values of mean front fitness over time for small *s* and other parameters close to values used in this paper, with relative difference 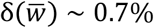 after 1000 generations. Unfortunately, this analytical model is known to underestimate actual mean front fitness, so the difference does not contribute to the accuracy of approximation of the actual mean front fitness. On the other hand, it differs from the formula in Peischl *et al*. (2013) quite drastically when *s* ≳ 0.03, *K* ≲ 10. Hereafter and in the results, we employ our derived formula (Eq. 2).

## 3. RESULTS

### 3.1. 1D

In one dimension, we have carried out 3 trials: one long simulation over 70 replicates, which is our main trial, designed to measure the long-term behaviour of fitness in range expansions (**1**), a trial concerning the critical value of the rate of deleterious mutations (**2**), and a trial employing a different approach to generating the offspring in the model (**3**). All trials are roughly one-directional, in the sense that individuals start in the leftmost demes of the habitat, and we measure the front as the (moving) populated outermost demes on the right.

We ran 70 simulation replicates, each with a burn-in phase and active phase of 10000 and 2500 generations respectively; as well as 70 replicates without the burn-in phase. Figure 2 shows the evolution of mean front fitness in time over 70 simulation replicates. Along with it, we have also run the program from Peischl *et al*. (2013) to simulate 10 and 20 replicates of the expansion with identical parameters (all of the parameters are given in Table 1). The rise after approximately 1930 generations of free expansion corresponds to the generation in which the population filled up the whole habitat, thus raising fitness due to efficient selection. For simulations with the burn-in phase was present, we also display the results of applying different normalisation methods (“onset mean normalisation” and “maximum normalisation” in Figure 2).

**FIGURE 2.**
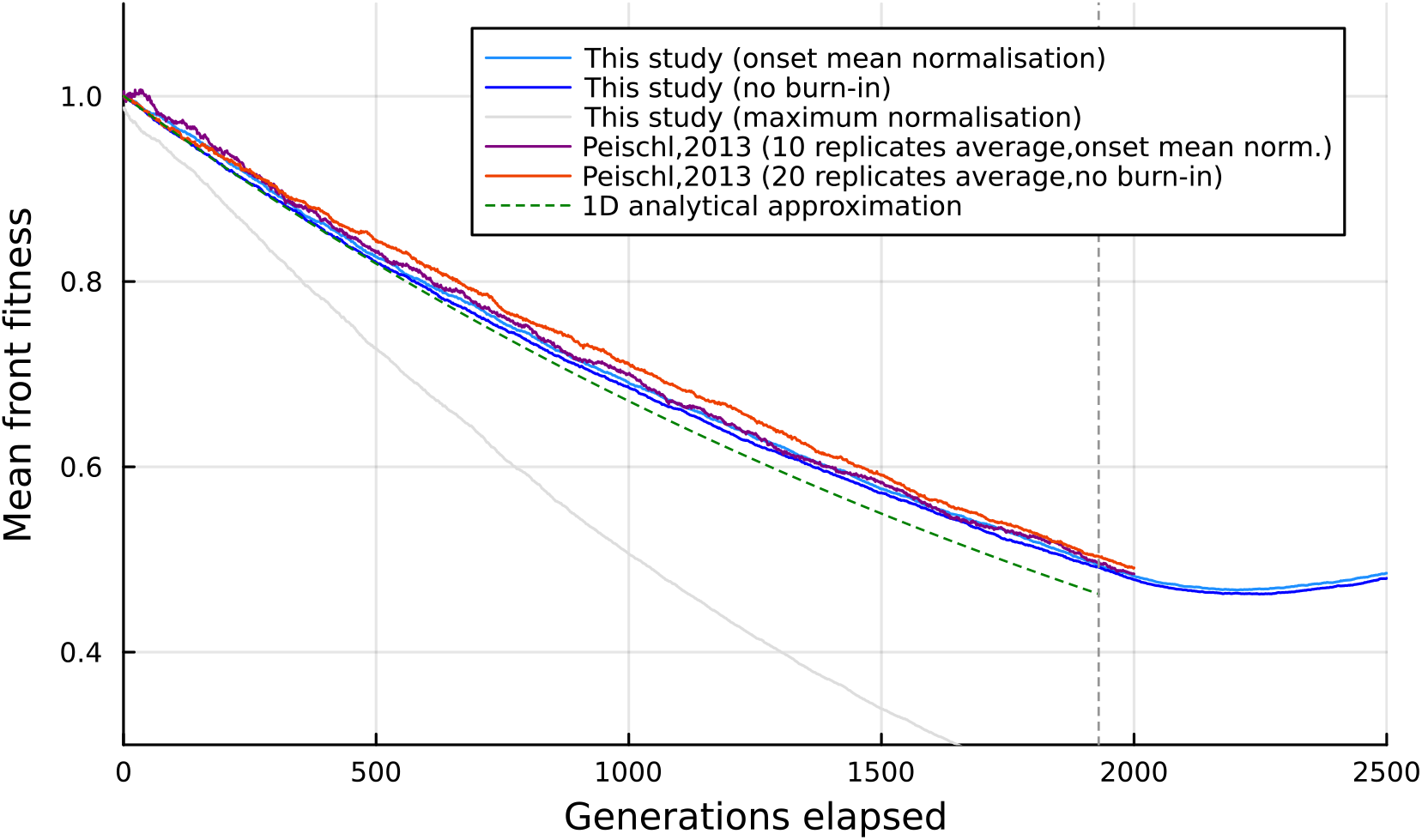
Temporal evolution of mean front fitness, as described in **1D 1)**. The grey vertical line signifies the generation when all demes in the space have been filled.

We have also performed a trial of the formula for the critical value of the rate of deleterious mutations, below which an expansion with the parameters used in **1)** (and given in Table 1) would always have increasing mean fitness at the front (Peischl *et al*., 2013). We calculated this value to be equal to ϕ_*c*_ ∼ 0.57. Thus we set the simulation parameters to the ones in **1)**, with the exception of ϕ = 0.45 and the simulation length: we run a simulation for 5000 burn-in generations and 1250 generations after the onset of the expansion, 10 times. The results confirm that our program gives an increasing trend of normalised mean front fitness during a range expansion, evenwith 10 simulation replicates (Figure S2).

To test out and compare the “first pair” and the “draw until pass” approaches that we have described in Chapter 2.1, we ran two sets of simulations under more extreme conditions. The results are shown in Figure S3. We can see that for mean front fitness with applied “onset mean normalisation”, the trend in both approaches is identical, and the difference in values is within 2%, excluding the rough fluctuations near the onset of the expansion (presumably due to a small sample of simulations).

### 3.2. 2D strip

In two dimensions, we likewise focused on developing simulation programs capable of confirming the results of prior research, concerning the evolution of mean fitness on the expansion front, as well as simulating and visualising the evolution of average fitness at every deme. Specifically, we developed two programs in 2D: expansion on a strip, and expansion on a disk. Below we talk about the main trial we carried out with the former program.

We created a program to simulate a range expansion roughly along an axis in two dimensions. We call this “2D strip” expansion, due to the habitat constituting a long rectangle, and give the parameters in Table 1. Individuals start in the leftmost demes of the habitat, and we measure the front as the (moving) populated outermost demes on the right. We ran 20 simulation replicates, each with a burn-in phase and an active phase of 1000 generations both; and the same number of replicates without the burn-in phase. Figure S4 (top) shows the evolution of mean front fitness in time for these results, averaged over replicates. We have also displayed the non-normalised mean front fitness for reference. Furthermore, we have displayed the results of applying different normalisation methods.

In addition, we have generated animations of a heatmap that shows the evolution of deme-average fitness throughout the whole expansion at every point in the habitat. Specifically, we generated heatmaps for deme-average fitness and normalised deme-average fitness. In this case, normalisation is such that the average fitness 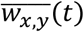 at each deme (*x, y* are space coordinates, *t* counts generations from the onset of the expansion; negative *t* is allowed in the presence of a burn-in phase) is divided by 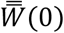, the average fitness over the whole habitat at the onset of the expansion. These animations are available in the “Data availability statement”. A snapshot of the heatmap of deme-average fitness is given in Figure S4 (bottom).

### 3.3. 2D disk

Next, we have simulated a range expansion on a 2D disk, which is bounded by the circle equation at a certain radius -- parameters are given in Table 1. Aside from this space boundary, the individual movement is not constrained. At the start, the founder individuals were put in the centre of the habitat. We have performed one trial, where we likewise found the evolution of mean front fitness in time, averaged over 20 simulation replicates. This result, with different normalisation methods, is shown in Figure 3. Similar to 2D strip, we have generated heatmaps that show the temporal evolution of (normalised) average fitness at every deme in the habitat, at every generation. This heatmap is available as an animation (see “Data availability statement”). A snapshot of the heatmap of deme-average fitness is given in Figure 3 (right).

**FIGURE 3.**
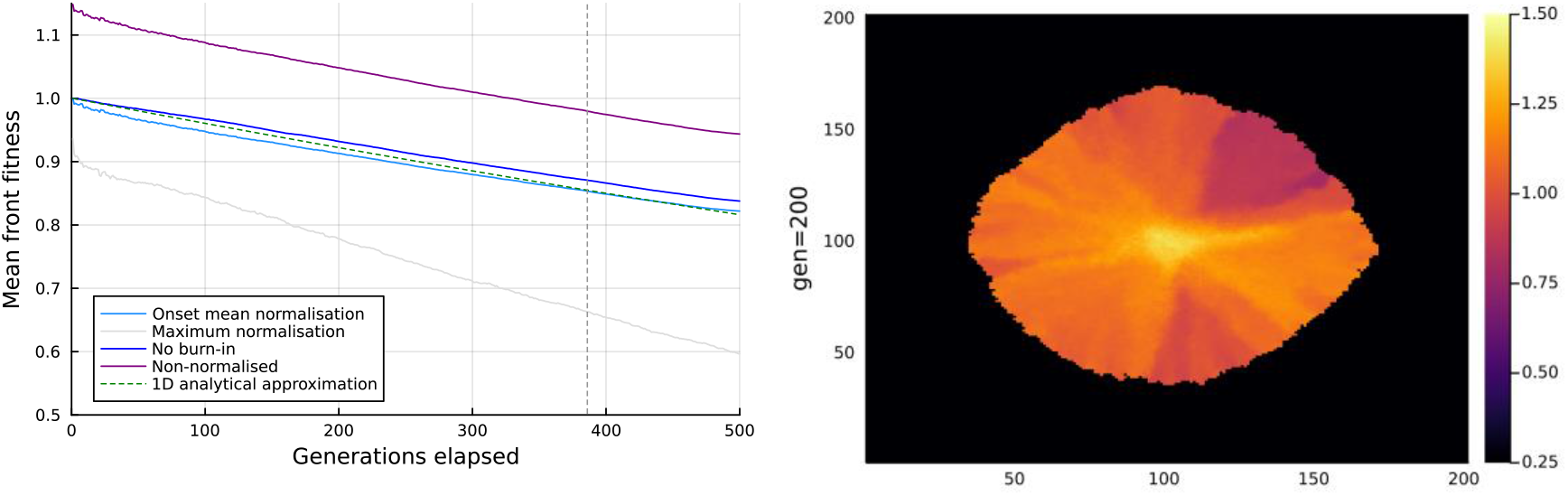
<u>Left</u>: temporal evolution of mean front fitness, as described in **2D disk**. The grey vertical line signifies the generation when all demes in the space have been filled. <u>Right</u>: a snapshot of the evolution of deme-average fitness (given by the colour) at every deme in this expansion, at generation 200.

### 3.4. 3D cylinder

The main goal of this study was to build a simulation tool that can produce data for range expansion in three dimensions, and analyse the evolution of mean front fitness consistently with the results from 1D and 2D. In particular, we developed two methods for 3D: expansion in a cylindrical habitat, and expansion in a sphere. Here, first, we describe the trial we carried out using the former program.

In this trial, we contained a population inside of a cylinder, that is constrained by a circle equation at the lateral surface and a barrier at both ends of the cylinder. This trial was roughly one-directional, meaning that the individuals started in the lowermost demes of the *z*-axis, and we measured the front as the (moving) populated highest demes in the direction of the *z* -axis. We ran 10 simulation replicates, each with a burn-in phase and an active phase of 1000 generations both; and the same number of simulation replicates without the burn-in phase. Figure 4 (left) shows the evolution of mean front fitness in time, averaged over 10 simulation replicates. As before, here we also display the non-normalised mean front fitness for reference and the results of applying different normalisation methods. We have also generated three-dimensional heatmaps of deme-average fitness, a snapshot of which can be seen in Figure 4 (right). The animations of these heatmaps are available in the “Data availability statement”. A typical slice image of the heatmap at height *z* =22 and generation 200 is presented in Figure S5 (left).

**FIGURE 4.**
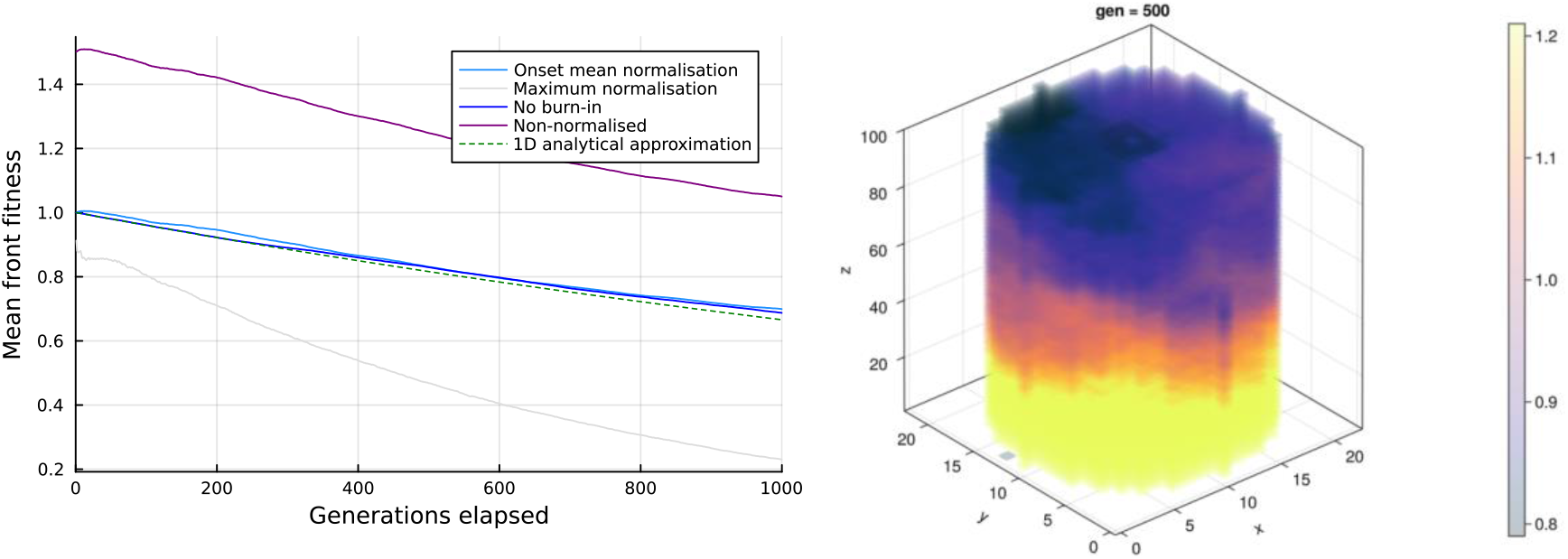
<u>Left:</u> temporal evolution of mean front fitness, as described in **3D cylinder**. <u>Right</u>: a snapshot of the evolution of deme-average fitness (given by the colour) at every deme in this expansion, at generation 500.

### 3.5. 3D sphere

Finally, we developed a simulation program for an expansion bounded by a 3D sphere of a constant size. At the start, the founder individuals were put in the centre of the habitat. We have performed one trial, where we likewise found the evolution of mean front fitness front in time, averaged over 10 simulation replicates. The result, with different normalisation methods, is shown in Figure 5. Similar to other trials, we have generated heatmaps for deme-average fitness at each generation. Sample animations of these heatmaps are available in the “Data availability statement”. A snapshot of the heatmap of deme-average fitness is given in Figure 5 (right). A typical slice image of the heatmap at height *z* = 18 and generation 642 is presented in Figure S5 (right).

**FIGURE 5.**
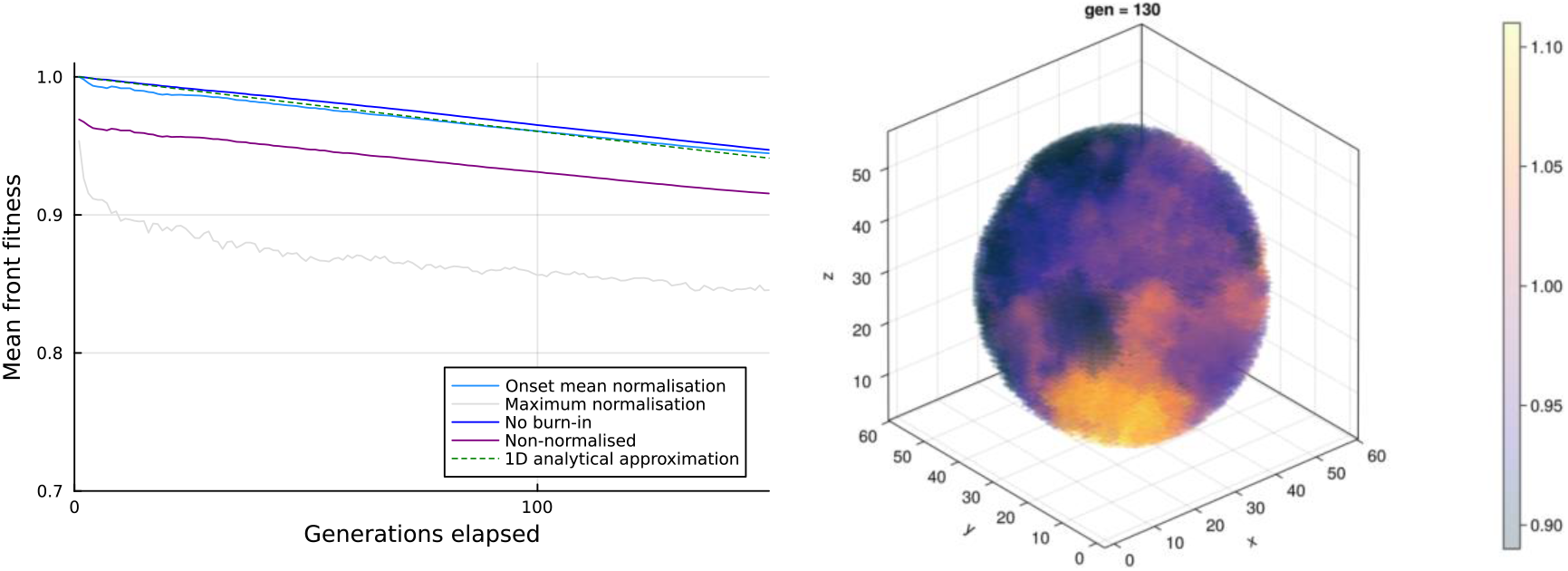
<u>Left:</u> temporal evolution of mean front fitness, as described in **3D sphere**. The grey vertical line signifies the generation when all demes in the space have been filled. <u>Right</u>: a snapshot of the evolution of mean deme fitness (given by the colour) at every deme in this expansion, at generation 130.

**FIGURE 6.**
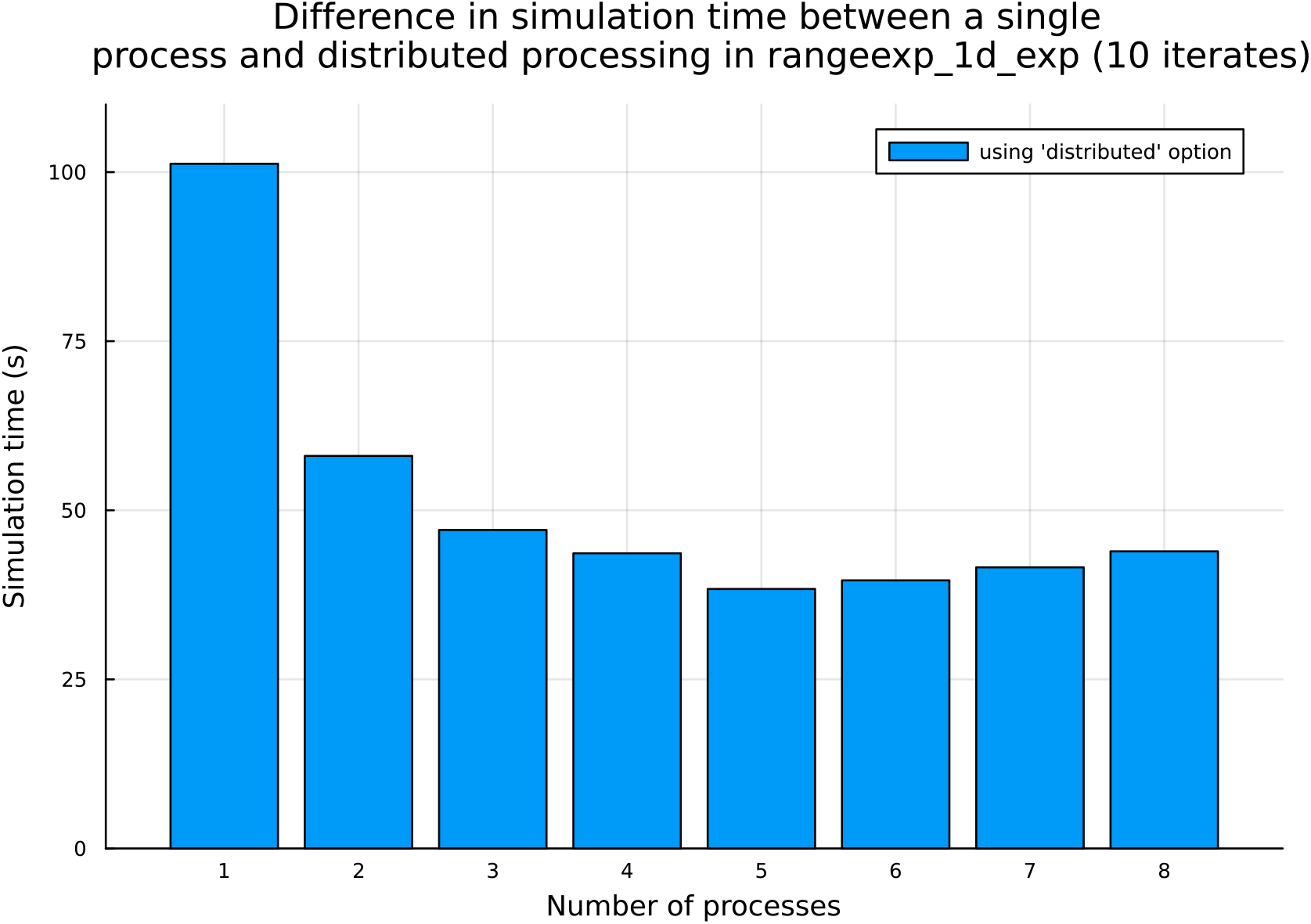
The results of a simulation time test done with a variable number of processes (from 1 to 10), to simulate 10 replicates of a range expansion in 10×100 deme habitat, for 100 (burn-in) + 200 generations. The method *rangeexp_1d_inf* from RESK was used.

## 4. DISCUSSION

### 4.1. 1D

Even though we have only used 10 and 20 replicates of simulation data generated with the program provided in previous research, we can see that simulations generated with our program yield close values and behaviour (relative error of normalised mean front fitness ≤ 1% everywhere expect the first 170 generations). The discrepancy between the analytical approximation and simulatedvalues grows visibly during longer expansions.

We have confirmed that the onset mean normalisation method yields values of mean front fitness for simulations with a burn-in phase closest in value to mean front fitness for simulations without a burn-in phase. Cf. the normalisation method where all demes are mapped to the interval [0, 1] every generation (“maximum normalisation” in Figure 2). The rapidly decreasing mean front fitness normalised via “maximum normalisation” can be used to measure the growing difference between the fitness in the core population and on the expansion front.

In trial **3**, we saw that the population cannot go extinct in range expansions in idealistic settings with soft selection (cf. range shifts, Gilbert *et al*. (2018)). As expected, in severe conditions, the mean front fitness in “first pair” populations was extremely low compared to “draw until pass” populations. However, when applying the “onset mean normalisation” method, these fitness values were almost equal throughout the expansion. Hence, we can say that onset mean normalisation preserves the trend well, and mean front fitness for these two offspring generation approaches differs only in scale of the absolute value.

### 4.2. 2D

Our 2D disk simulations show that, in an expansion where the population is free to move in 4 directions (orthogonally), segregation patterns resembling spokes or circular sectors emerge practically from the start of the expansion (Figure 3). This behaviour has been observed extensively in bacterial mixtures with only a single mutant allele (Hallatschek and Nelson, 2010; Korolev *et al*., 2010). Moreover, the behaviour was recently confirmed in originally monospecific populations partially or fully developing specialist subpopulations during the expansion (Li and Kardar, 2022). Our results show that this binary segregation behaviour (i.e. with zones where the mutant allele frequency is strictly 0 or 1) changes into a picture with smooth transitions between predominantly mutant and predominantly non-mutant zones in expansions of populations with multiple mutation types. This is speculated to be a blend of separate binary segregation pictures against each of those mutations (no matter received before or during the expansion). This was shown in part in Excoffier *et al*., 2008. The simple model used in this study can be approximated by a model with co-dominant mutations for low *s*, and it should be noted that for dominant or recessive mutations this behaviour might be different. It might also be different when long-range migration is allowed: in their simulations, Paulose and Hallatschek (2020) found that the possibility for long-range dispersal could make the segregating pattern above disappear.

On the other hand, in our 2D strip trial shown in Figure S4, large-scale spatial deviations from the monotonous decrease in fitness with distance from the core population were not present in the narrow space we have set up. We’ve run separate trials to confirm that this is only the case with narrow expansions that approach an axial expansion such as the 1D expansion, and is also dependent on the simulation parameters such as the selection coefficient and migration rate. Cf. other simulations with wide rectangles (0 ≤ *y* ≤ 20) (Peischl *et al*., 2015b).

### 4.3. 3D

This is the first time that range expansions of this type were simulated in 3D. The deme-average fitness in**3D cylinder** (Fig. 8, right) was found to exhibit patterns of smooth combination of binary segregation pictures, similar to the those observed in **2D disk** (Fig. 7, right), but with more variation in shape, nomore resembling spokes or circular sectors. This pertains to the concurrent variation in the lateral *xy* plane with the variation along the *z* -coordinate. The effect from this joint variation seems greater that the effects from separate respective variations. The combination of radial and axial variations here provides a more extreme fitness distribution, where any slight deviation from the central expansion axis has a chance to result in a drastic falloff in fitness. This suggests that, aside from the monotonous decrease in fitness proportional to the distance from the origin, colonisers in 3D space need to take into account the dramatic variation in fitness when straying from the central axis.

On the other hand, the mean front fitness can still be approximated fairly well using the analytical formula for one-directional 1D expansions. This is also the case for the spherical expansion (Fig. 9). Moreover, in all of the expansion scenarios in 2D and 3D, meanfront fitness could still be well approximated with the analytical formula for simple one-directional 1D expansions. We assume this follows from the fact that, for each deme at the front, the probability to colonise a further deme is approximately equal to the probability to remain in the present population. However, a thing to note is the sudden drop in mean front fitness just after the onset of the expansion in simulations with a burn-in phase. This is a flaw of the burn-in phase, and could be explained by the fact that the population at the time of the onset experiences the lifting of a migration restriction, and this allows it to migrate and undergo the biggest loss of genetic diversity per deme, since the population is still comparatively small.

Genetic drift was shown to be a factor in heterozygosity gradients in a population (Liu *et al*., 2006), and in future studies, it would be of value to analyse the distribution of heterozygous loci and calculate the temporal change in expansion-front heterozygosity in different dimensions. We have already developed the methods for such simulations that use a finite-site model with dominance – these methods are available in RESK. Upon modifying our programs to a finite number of loci (1000 and more), the calculations turned out to be computationally intensive in multiple dimensions, increasing the simulation time severalfold compared to the infinite-sites finite-region model in 3D. We would like to explore this model, using more powerful hardware in a future study, as well as investigate the difference between soft and hard selection. Nonetheless, the infinite-site finite-region model explored in this study proved to be an efficient tool for studying the general properties of expansion load.

## 5. CONCLUSION

A program kit was developed for efficiently simulating range expansions in different habitats in 1, 2 and 3 dimensions. Using them, the mean fitness of expansions in three dimensions was successfully simulated for the first time. Different normalisation methods were investigated. “ Maximum normalisation” was found to be useful to measure the growing difference between the fitness in the core population and on the expansion front. In radial 2D (disk) expansions, deme-average fitness of a population that underwent multiple mutations was found to show a smooth combination of binary segregation pictures against each of those mutations. This pattern was also observed as a component in the deme-average fitness in 3D expansions. Overall, complex fluctuations in deme-average fitness, different from those in radial 2D expansions, were found in 3D expansions. Axial 3D (cylinder) expansions were found to have the highest fitness along the axis of the expansion.

## Author contributions

Conceptualisation and providing the calculation environment: Gen Tamiya.

Simulation software development: Hikaru Matsuoka and Yurii Kotsar.

Planning and creating example simulations used in the study: Yurii Kotsar.

Manuscript: Yurii Kotsar.

Manuscript revision and corrections: Hikaru Matsuoka and Gen Tamiya.

## Data availability statement

We have uploaded the program kit, some simulated result data, as well as animations used in the study to: *https://github.com/HartreeY/RESK*. Thes program kit consists of the core file (*resk*.*jl*) with all the simulation methods, including *rangeexp_inf* – the one used in this study; an optional graphing methods file (*reskplots*.*jl*); a file for defining default parameters (*consts*.*jl*); and the initialisation file that installs the requirements for this program kit (*init*.*jl*). We have also included examples in the *study* folder that correspond to trials in this study. In it, we have also uploaded animations of the heatmaps for results of 2D and 3D simulations to the *animations* folder.

To generate the data for comparison with Peischl *et al*. (2013), we used the following program linked in their paper: *https://github.com/CMPG/ADMRE/tree/master*.

## APPENDIX A APPENDIX A. Floating point error

Due to operating memory limitations, we used Float32 arrays to store fitness values. Since the fitness of foundersexponentially influences the fitness of next generations, we designed a test for cumulative errors in mean front fitness values.

We performed the test both for 1D and 3D. In 1D, we ran 15 simulation replicates with the default parameters given in Table 1, however only for 1000 burn-in generations and 1000 generations after onset. These were run both by a program which employs Float32, and one which employs Float64. For every simulation, we synchronised the random seeds given to both programs. We calculated the average over 15 replicates of the mean front population and mean front fitness for each generation, for outputs of both of the programs. We then analysed the relative difference between the averages in the two programs. The difference in average population was negligible, with relative error ∼0.3%. The difference in average deme fitness was likewise unimportant with the value < 0.3%.

In 3D, we ran 10 simulation replicates for both programs with the default parameters given in Table 1, but setting 100 burn-in generations and 600 generations after onset. Both differences had a maximum at the last generation inthe simulation, indicating that this could be a cumulative floating-point error. The maximum of the relative difference in mean front population was ∼0.6%, while the maximum of the relative difference in mean front fitness was also ∼0.6%.

It goes without saying, that this is a comparison between Float32 and Float64, and not between Float32 and anideal floating-point datatype. Still, this error component was found not likely to influence the results of the tests described above in a significant way.

## APPENDIX B APPENDIX B. Parallel processing

For our simulation kit, we have included the possibility to run parallel simulations, to generate multiple replicates in one go. The results in this study were generated in parallel. In Figure 6, we show a simple test, which we have also included in *https://github.com/HartreeY/RESK/programs/examples/distributed_2.ipynb*. The test compares computation times to run 10 replicates of the same simulation on a variable number of parallel processes: from 1 to 8. The test was performed on a machine with 8 CPUs. Figure 6 shows that even with only 2 processes, the performance is boosted almost twofold, and additional processes further decrease the time in a diminishing manner (although there is a reverse effect due to the small amount of CPUs). In scientific research, having data from a large number of replicates is invaluable, and the built-in option of parallel simulations should add to the efficiency of generating data for studies that rely on simulations.

