## Supplemental Tables for "The evolution of fitness during range expansions in different dimensions"

**Table of Contents:**

| **Figure S1** | Page 1 |
| --- | --- |
| **Figure S2** | Page 2 |
| **Figure S3** | Page 2 |
| **Figure S4** | Page 3 |
| **Figure S5** | Page 3 |

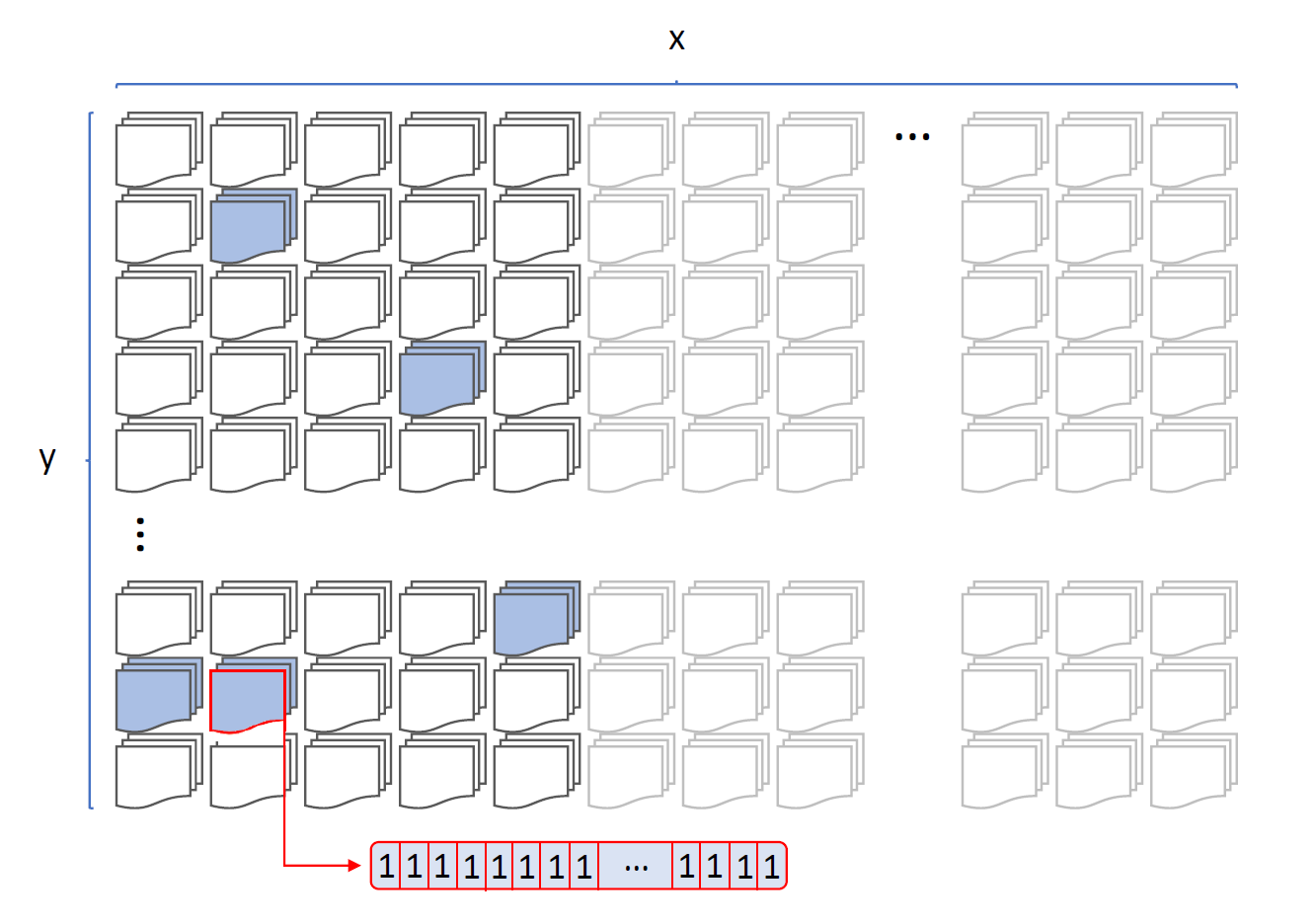

**FIGURE S1.** The habitat and its arrays of individuals, in the example of a 2D simulation. The x and y axes represent habitat dimensions and are given in Table 1 for specific simulations. Demes start out empty, except for *b* random demes highlighted in blue. Such demes house 100 individuals: as an example, one is highlighted in red. Their fitness values are all set to 1 at start. The faded out grey arrays/demes are unreachable during the burn-in phase.

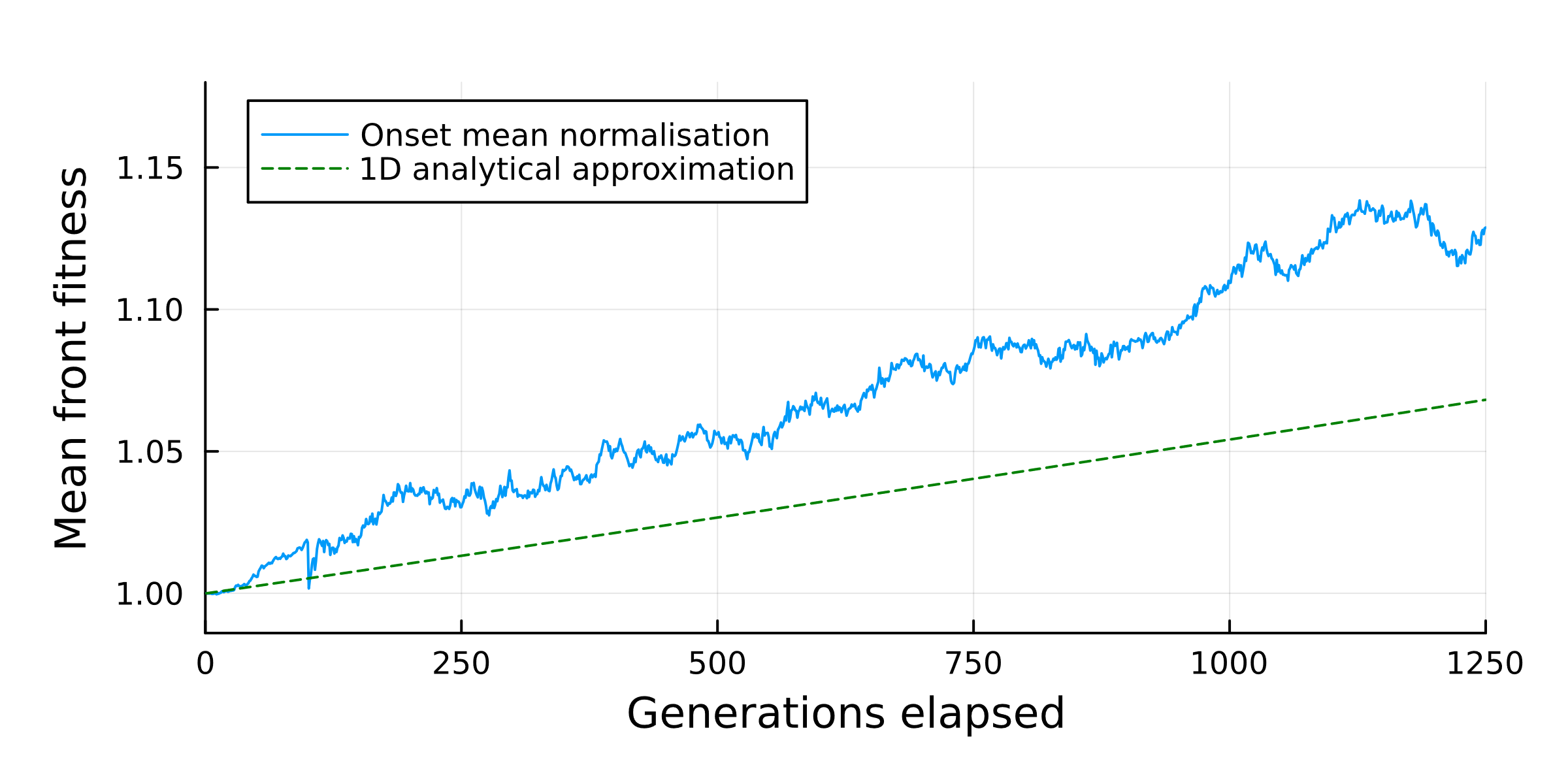

**FIGURE S2.** Temporal evolution of mean front fitness of a trial described in **1D 2)**.

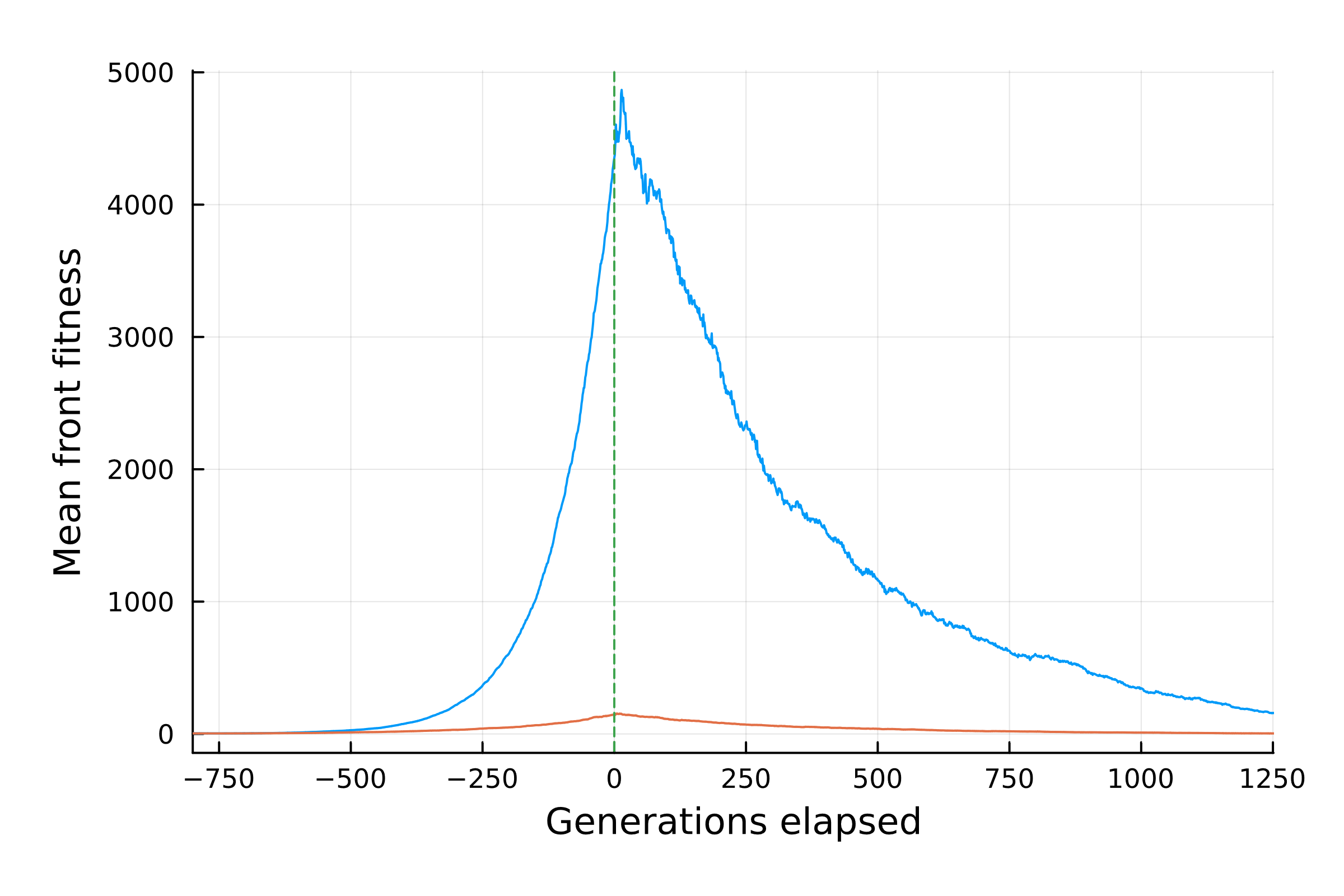

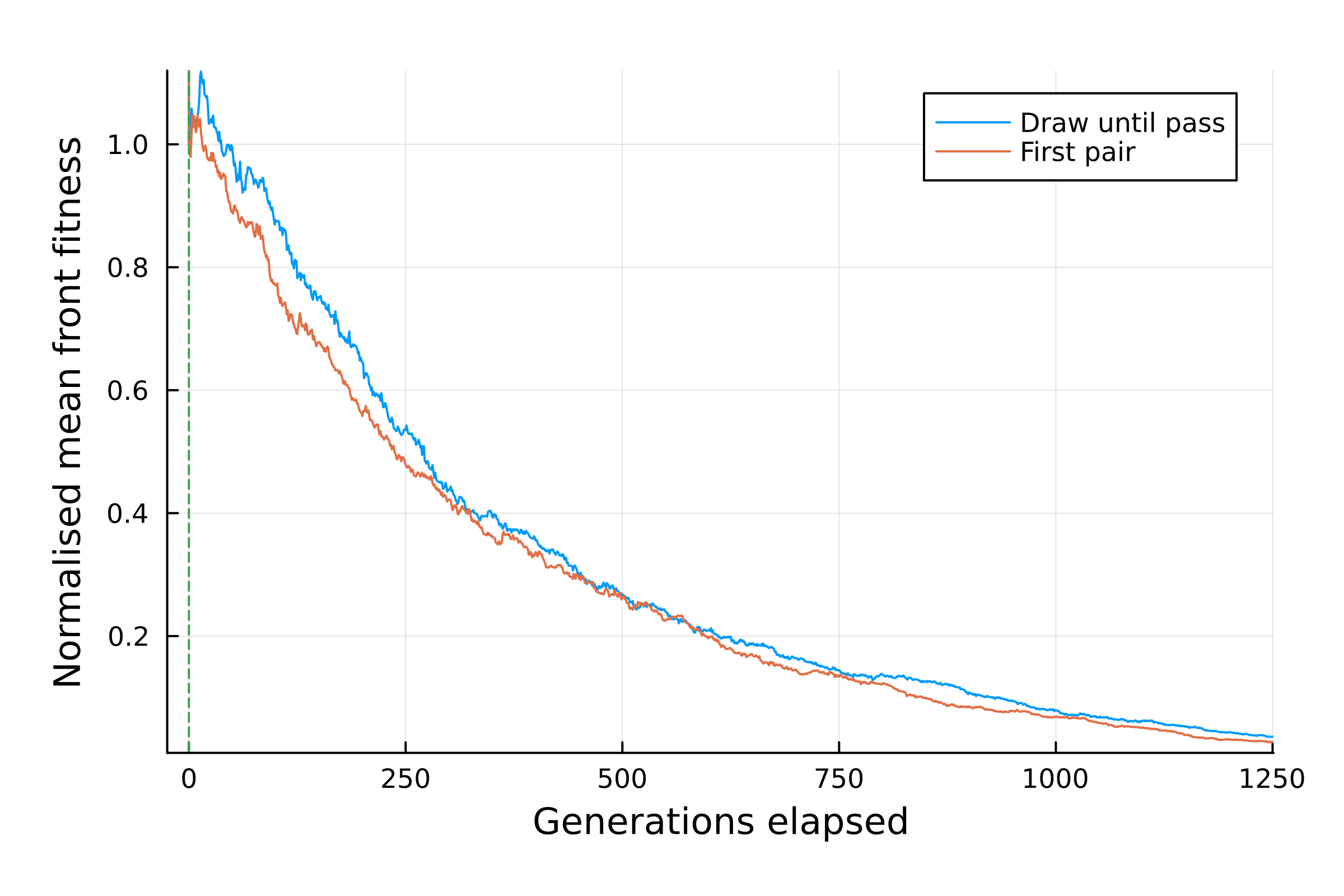

**FIGURE S3.** Left: temporal evolution of mean front fitness as described in **1D 3)**. The green vertical line signifies the onset of the expansion; generations to the left of it constitute the burn-in phase. Right: the main expansion phase zoomed in, and mean front fitness further normalised according to the "onset mean normalisation" method.

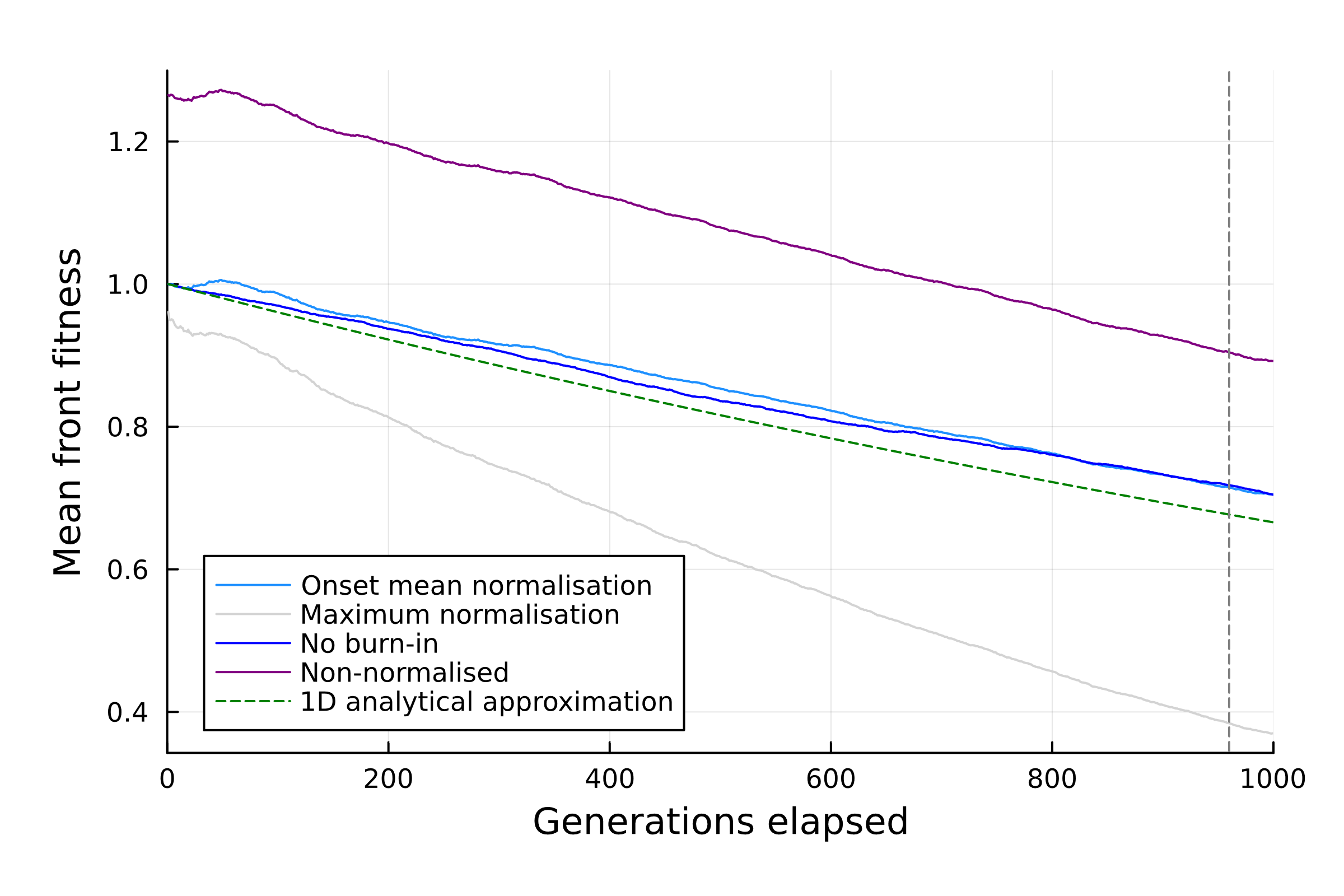

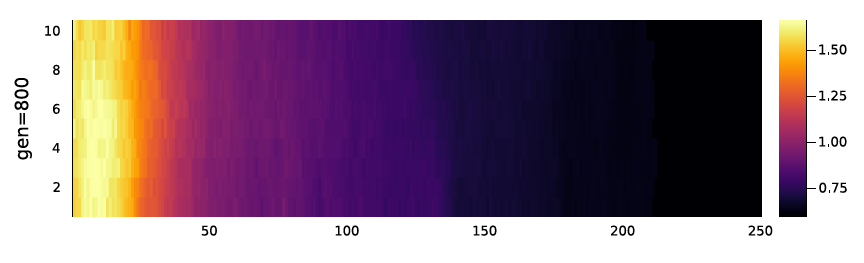

**FIGURE S4.** Top: temporal evolution of mean front fitness, as described in **2D strip**. The grey vertical line signifies the generation when all demes in the space have been filled. Bottom: a snapshot of the evolution of mean deme fitness (given by the colour) at every deme in this expansion, at generation 800.

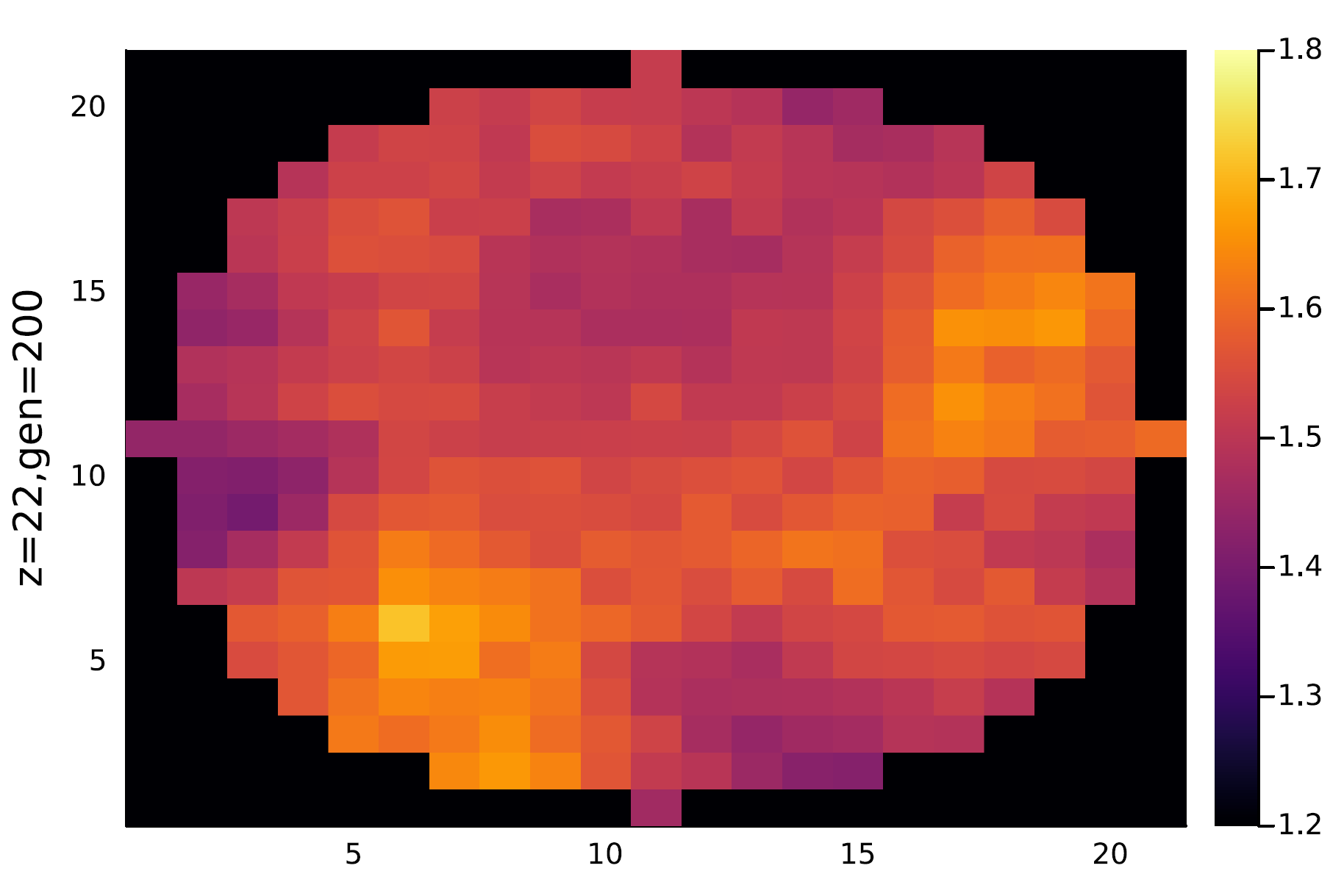

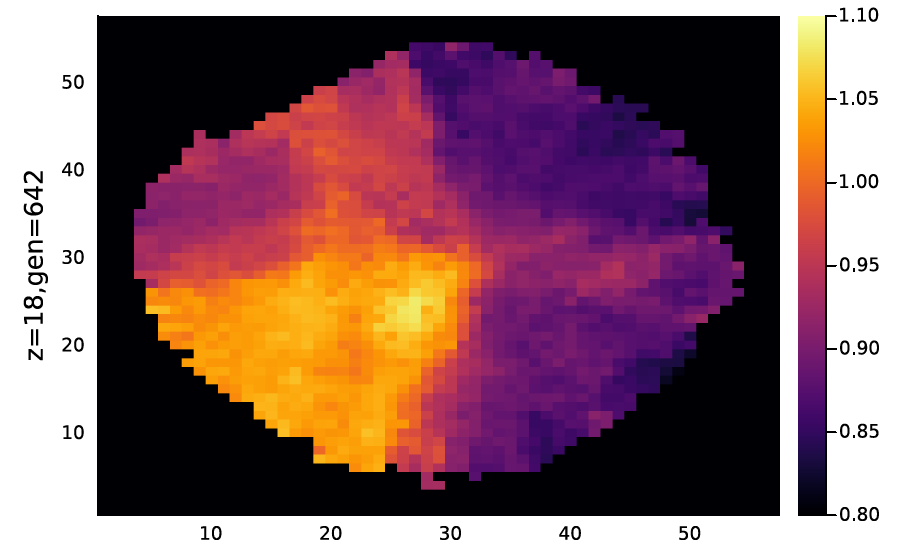

**FIGURE S5.** Left: a cross-section at height $z=22$, at generation 200 after the onset of an expansion described in **3D cylinder**. Right: a cross-section at height $z=18$, at generation 642 after the onset of an expansion described in **3D sphere**. In both graphs, the heatmap values show deme-average fitness.
